# The origin and evolution of a distinct mechanism of transcription initiation in yeasts

**DOI:** 10.1101/2020.04.04.025502

**Authors:** Zhaolian Lu, Zhenguo Lin

## Abstract

The molecular process of transcription by RNA Polymerase II is highly conserved among eukaryotes (“classic model”). Intriguingly, a distinct way of locating transcription start sites (TSSs) was found in a budding yeast *Saccharomyces cerevisiae* (“scanning model”). The origin of the “scanning model” and its underlying genetic mechanisms remain unsolved. Herein, we applied genomic approaches to address these questions. We first identified TSSs at a single-nucleotide resolution for 12 yeast species using the nAnT-iCAGE technique, which significantly improved the annotations of these genomes by providing accurate 5’boundaries of protein-coding genes. We then infer the initiation mechanism of a species based on its TSS maps and genome sequences. We found that the “scanning model” had originated after the split of *Yarrowia lipolytica* and the rest of budding yeasts. An adenine-rich region immediately upstream of TSS had appeared during the evolution of the “scanning model” species, which might facilitate TSS selection in these species. Both initiation mechanisms share a strong preference for pyrimidine-purine dinucleotides surrounding the TSS. Our results suggested that the purine is required for accurately recruiting the first nucleotide, increasing the chance of being capped during mRNA maturation, which is critical for efficient translation initiation. Based on our findings, we proposed a model of TSS selection for the “scanning model” species. Besides, our study also demonstrated that the intrinsic sequence feature primarily determines the distribution of initiation activities within a core promoter (core promoter shape).

## INTRODUCTION

Transcription of protein-coding genes by RNA polymerase II (Pol II) is an essential process in the “central dogma” that converts genetic codes in DNA into functional products. A crucial step of transcriptional regulation occurs at transcription initiation, as it determines not only the number of transcripts produced but also the locations of transcription start sites (TSSs). Therefore, transcription initiation has been a focus of many studies of gene regulation (Roeder 1996). Genome-wide studies in various eukaryotic organisms revealed that transcription initiation is highly pervasive and dynamic (Carninci et al. 2005; Carninci et al. 2006; Hoskins et al. 2011; Encode Project Consortium 2012; Lu and Lin 2019). Alternative usage of TSS is prevalent in response to environmental cues, and it is usually associated with gene differential expression (Lu and Lin 2019). It was shown that transcript isoforms produced by alternative TSSs have different translation efficiency (Cheng et al. 2018). From an evolutionary perspective, changes in TSSs were found to be associated with divergence of gene expression patterns and phenotypic traits (Lin and Li 2012).

Most studies of Pol II transcription machinery were performed with promoters with a TATA box (Patikoglou et al. 1999), which is the first core promoter element identified (Smale and Kadonaga 2003). The process of transcription initiation from TATA box-containing promoters is highly conserved from archaea to eukaryotes. In brief, general transcription factors (GTFs), including TFIIB and the TFIID subunit of TATA box binding protein TBP, recruit Pol II to form a preinitiation complex (PIC) that allows Pol II to reach TSS directly (Bernard et al. 2010; Li et al. 2015; Blombach et al. 2016). Therefore, transcription is initiated at ~30 base pairs (bp) downstream of the TATA box, termed as “classic model” herein. Intriguingly, a distinct mechanism of transcription initiation has been observed in a budding yeast *Saccharomyces cerevisiae* (Choi et al. 2002; Hahn and Young 2011). Specifically, the PIC in *S. cerevisiae* performs a scanning process to seek favorable TSSs and initiates transcription mainly from 60 to 120 bp downstream, denoted as the “scanning model” (Giardina and Lis 1993; Kuehner and Brow 2006; Fishburn and Hahn 2012). A recent study suggested that the “scanning model” is also used in promoters without a TATA box in *S. cerevisiae* (Qiu et al. 2019), suggesting it is a genome-wide transcriptional initiation mechanism. A fission yeast *Schizosaccharomyces pombe* demonstrates a similar initiation pattern to “classic model” species in TATA-containing promoters (Choi et al. 2002). Thus, the divergence of transcription initiation mechanisms might have occurred after the split of fission yeasts from budding yeasts. As the two lineages diverged over 500 million years ago (Rhind et al. 2011), the more accurate timing of the origin of the “scanning model”, as well as its underlying genetic basis, has yet to be determined.

Studying the evolution of transcription initiation mechanisms could provide a better understanding of the molecular mechanisms underlying how the PIC identifies TSS. For instance, in both “classic model” and “scanning model” species, transcription is mostly initiated from a purine (TSS or position +1) at the sense strand, with a pyrimidine immediately upstream of it (position-1), which is called pyrimidine-purine (PyPu) dinucleotide (Carninci et al. 2005; Hoskins et al. 2011). It has been shown that the −1 pyrimidine facilitates the stacking of the first NTP by Pol II (Zhang et al. 2014). However, it remains unclear why purine is strongly preferred as the first recruited nucleotide during transcription. Besides, an adenine was found at eight bp upstream of most TSSs (abbreviated as −8A hereafter) in *S. cerevisiae* (Zhang and Dietrich 2005; Lu and Lin 2019). A structural study showed that the −8A is recognized by the B-reader helix of TFIIB, which is important for TSS selection (Kostrewa et al. 2009). And yet, whether the preference of −8A is also present in other “scanning model” species is not known, which is necessary to better understand its functional importance in these species.

Transcription within a core promoter is commonly initiated from a cluster of nearby TSSs, instead of a single TSS (Carninci et al. 2005). The distribution of transcription activities among TSSs in a core promoter varies substantially, forming different core promoter shape. Without an explicit cutoff, for simplicity, core promoters were generally divided into classes of “sharp” and “broad” (Fischer et al. 2000; Carninci et al. 2006; Sandelin et al. 2007). In mammals, broad core promoters are mostly present in ubiquitously expressed genes, while sharp core promoters are often associated with tissue-specific expressed genes (Carninci et al. 2006). By examining the TSS maps for 81 lines of *D. melanogaster*, Shor et al. identified thousands of genetic variants that could impact the transcription level and core promoter shape (Schor et al. 2017). However, the determinant factors of core promoter shape are not entirely understood.

With an aim to address the above questions, here we obtained and analyzed TSS maps at a single-nucleotide resolution for 12 yeast species, including ten budding yeasts and two fission yeasts. Based on in-depth interrogation of these TSS maps, we have pinpointed the origin of the “scanning model”, and inferred key genetic innovations associated with its evolutionary process. Our results also demonstrate the role of core promote sequences in transcription initiation and the genetic determinants of core promoter shape. Furthermore, we provided a plausible explanation for the evolutionary divergence in the length of 5’untraslated regions (5’UTR) in yeasts. These findings improved our understandings of the evolutionary divergence of transcription initiation mechanisms and the functional roles of sequencing elements in the key process of the “central dogma”.

## RESULTS

### Evolutionary dynamics of transcription initiation landscapes in yeasts

We generated high-resolution TSS maps for ten budding yeast species and two fission yeast species, including *S. cerevisiae*, *Sch. pombe*, and other important species, with estimated divergence times ranging from 4 to over 500 million years (Fig. 1A; Supplemental Table S1). These TSS maps were obtained using the no-amplification non-tagging cap analysis of gene expression (nAnT-iCAGE) technique (Murata et al. 2014). A total number of 838 million CAGE tags from the 12 yeast species were produced (Supplemental Table S2). The average overall mapping rate is 88.28%, and 617 million CAGE tags were uniquely mapped to the 12 genomes (Supplemental Table S2). We applied the Poisson model to remove TSSs that are likely due to stochastic transcription activities or transcription from non-*bona fide* core promoters (see Materials and Methods). On average, each species uses 286,433 TSSs grown in rich medium (Fig. 1A), supporting the pervasive nature of transcription initiation in yeasts, given their small genome size (~12M) and gene numbers (~5,000-6,000).

We developed a peak-based clustering (Peakclu) method to identify TSS clusters (TCs), representing core promoters (see Materials and Methods). We assigned TCs to Pol II transcribed genes as their core promoters based on their position proximity (see Materials and Methods). We identified core promoters for 83.72% protein-coding genes for these species (ranging from 4,571 genes in *Sch. japonicas* to 5,348 genes in *S. cerevisiae*). We defined the representative TSS of a protein-coding gene as the TSS with the highest CAGE signal in its promoter region. Our core promoter and TSS data improve the annotations of these genomes by providing 5’ boundaries for most genes at a single-nucleotide resolution. The updated genome annotations of these yeast species were provided as Supplemental Dataset S1-S12.

**Figure 1:**
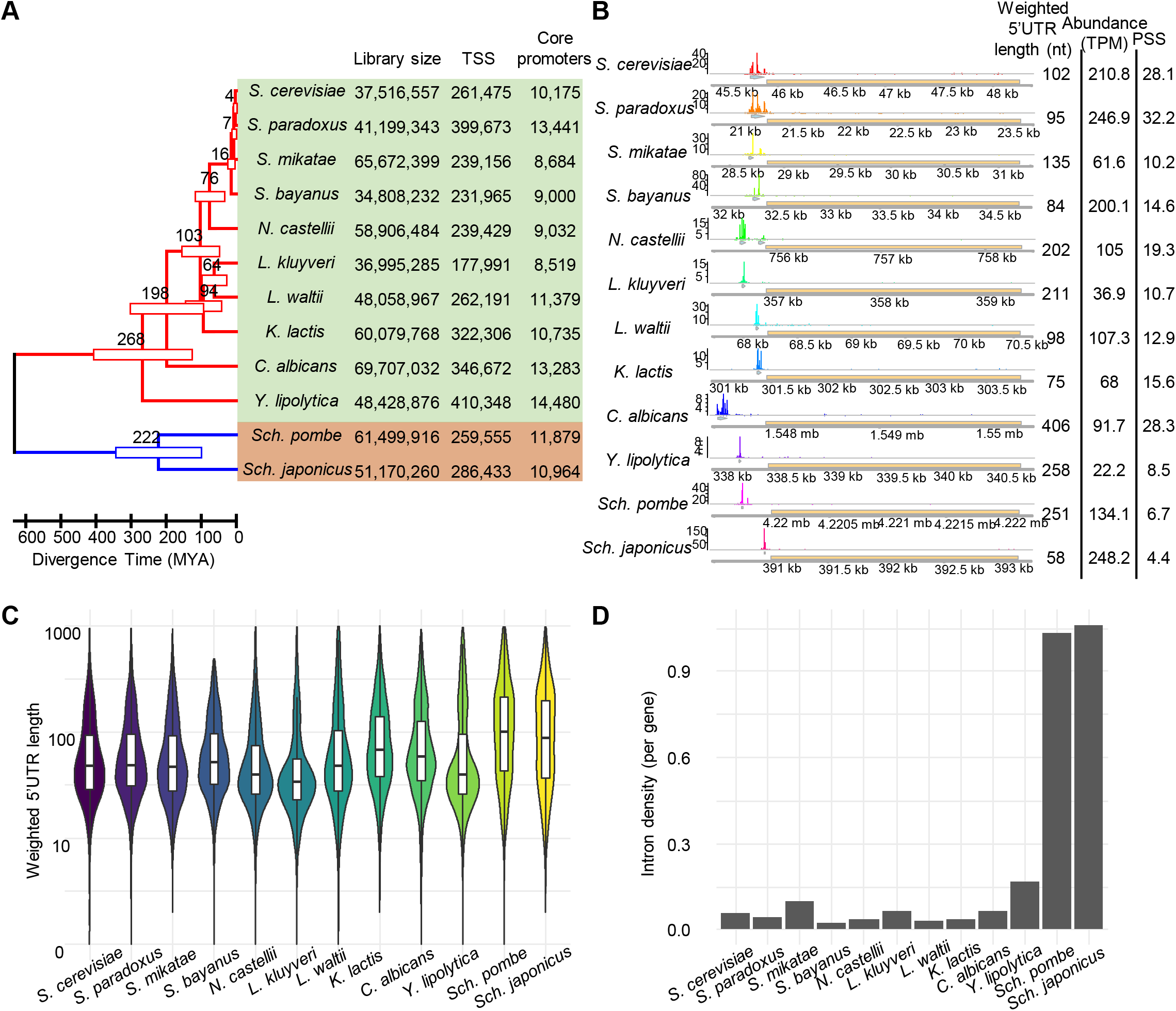
Genome-wide identification of TSSs in yeasts. (A) Phylogenetic relationships of the 12 yeast species examined in this study. The phylogenetic tree was inferred based on the largest subunit of Pol II RPB2 protein sequences by Maximum Likelihood method. The table on the right side includes library size, numbers of TSSs, and core promoters identified in each species. The full species names are provided in Table S1. (B) An example of TSS maps of 12 orthologous genes of *FLC2*. In each species, the top track illustrates the distributions of TSS signals. The second track (light green arrow) represents the core promoter region. The third track (yellow box) shows the locations of gene coding regions. The bottom track displays genome coordinates. (C) Violin plot shows the distribution of 5’UTR lengths in each species. (D) Numbers of intron per gene in each of the 12 genomes.

To better characterize the evolutionary patterns of core promoters, we delineated protein-coding genes of the 12 yeast species into 6,614 orthologous groups by OrthoDB (Kriventseva et al. 2019) (Supplemental Dataset S13). We found that core promoters among orthologous genes tend to have distinct features, including transcription activities, 5’UTR length, and core promoter shape, as illustrated by using the orthologous group of *FLC2* as an example (Fig. 1B). These features are most similar between the closely related species, such as *S. cerevisiae* and *S. paradoxus*, which have diverged ~4 million years ago. Whereas, the similarities of these features reduced with increases in divergence times. For example, larger differences can be observed between *S. cerevisiae* and its second-most related species *S. mikatae*, suggesting that these features could be related to genetic factors. However, these differences are not positively correlated with divergence times on a broader scale, probably because changes in these features were not directional, or they might have diverged at a much higher rate than genomic sequences.

One of the most significant differences related to TSSs between budding yeasts and fission yeasts is the 5’UTR length. The budding yeasts have a shorter median length of 5’UTR than that in fission yeasts (Fig. 1C; Supplemental Dataset S14). 5’UTR length in fission yeasts is more similar to higher eukaryotes, such as tomato (106 nt) and cow (111 nt) (Leppek et al. 2018). We speculated that the shortening of 5’UTR length in budding yeasts might be due to the massive loss of intron in their common ancestor. Budding yeasts have a significantly lower intron density than fission yeasts (Fig. 1D). As exonization from introns in 5’UTR regions has occurred (Hooks et al. 2014), elongation of 5’UTR by exonization of introns is more likely to occur in the intron-rich genomes than in intron-depleted genomes. We speculated that the massive loss of introns in budding yeast genomes largely eliminates the possibility of 5’UTR elongation through the exonization process.

In each examined species, 5’UTR lengths vary dramatically among individual genes. We measured the coefficient of variation (CV) of 5’UTR lengths among orthologous genes of each KEGG pathway to quantify their evolutionary divergence. We found that KEGG groups with the most divergent 5’UTR lengths are mostly related to metabolism pathways, such as ether lipid metabolism and riboflavin metabolism (Supplemental Fig. S1A). The median lengths of 5’UTR among genes in the riboflavin metabolism pathway range from 21 nucleotides (nt) to 294 nt in these species (Supplemental Fig. S1B). In contrast, the KEGG groups with the most conserved 5’UTR length are enriched in the essential cellular function pathways, such as ribosome and RNA transport (median 5’UTR lengths ranges from 37 to 63 nt, Supplemental Fig. S1C). These results support that 5’UTR length, which is primarily determined by the location of TSSs, is related to gene functions and expression profiles (Lin and Li 2012), although its underlying mechanism remains to be further investigated.

### The “scanning model” emerged during the evolution of budding yeasts

A distinct feature between the “classic model” and “scanning model” is the distribution pattern of TATA box relative to the TSS. Therefore, we used this feature to infer the transcription initiation mechanism in a species. We searched for TATA box motifs using its consensus sequence TATAWAWR (Basehoar et al. 2004; Rhee and Pugh 2012) in the promoter regions in each species examined. We found a binary pattern of TATA box positioning among the 12 species (Fig. 2). In one group, including a known “classic model” species *Sch. pombe*, as well as the other fission yeast examined *Sch. japonicas* and a budding yeast *Yarrowia lipolytica*, TATA box motifs are well-positioned at ~30bp upstream of TSS, supporting that they all belong to the “classic model”. In the other group, which includes a known “scanning model” species *S. cerevisiae*, and all other budding yeasts but *Y. lipolytica*, TATA boxes are mainly distributed in a broad range from 60bp to 120 bp upstream of TSSs, suggesting that these species use the “scanning model” for transcription initiation (Fig. 2). *Y. lipolytica* is the earliest branching species among budding yeasts examined, which had diverged about 200 million years ago. These results support that the origin of the “scanning model” had occurred after the divergence of *Y. lipolytica* during the evolution of budding yeasts.

In “classic model” species, transcription from TATA-containing core promoters tends to be initiated from a narrow range of TSSs, so they have a sharper shape of core promoter than those TATA-less promoters (Carninci et al. 2006). We calculated the core promoter shape score (PSS) for both classes of core promoters in each examined species (see Materials and Methods). Our study of core promoter shape in these species led to two major findings. First, in TATA box-containing promoters, the PSS values in the three “classic model” species are significantly lower (sharper) than the nine “scanning model” species (Fig.2). Such a difference is probably due to different ways of locating TSSs between the two classes of species. In the “scanning model” species, the PIC scans DNA sequences downstream of a TATA box to select favorable TSSs instead of from a fix distance, leading to a broader distributed TSSs. These findings also support that our inference of transcription initiation mechanisms based on distributions of the TATA box is robust. Second, the PSS values of TATA box-containing promoters are significantly lower than TATA-less ones in “classic model” species, but such differences are absent in “scanning model” species (Supplemental Fig. S2). This observation suggests that the scanning mechanism is likely used for promoters with and without a TATA box in “scanning model” species, which is consistent with the results in a separate study (Qiu et al. 2019). Therefore, our findings based on core promoter shape further support that the “scanning model” originated after the split of *Y. lipolytica* during the evolution of budding yeasts.

**Figure 2:**
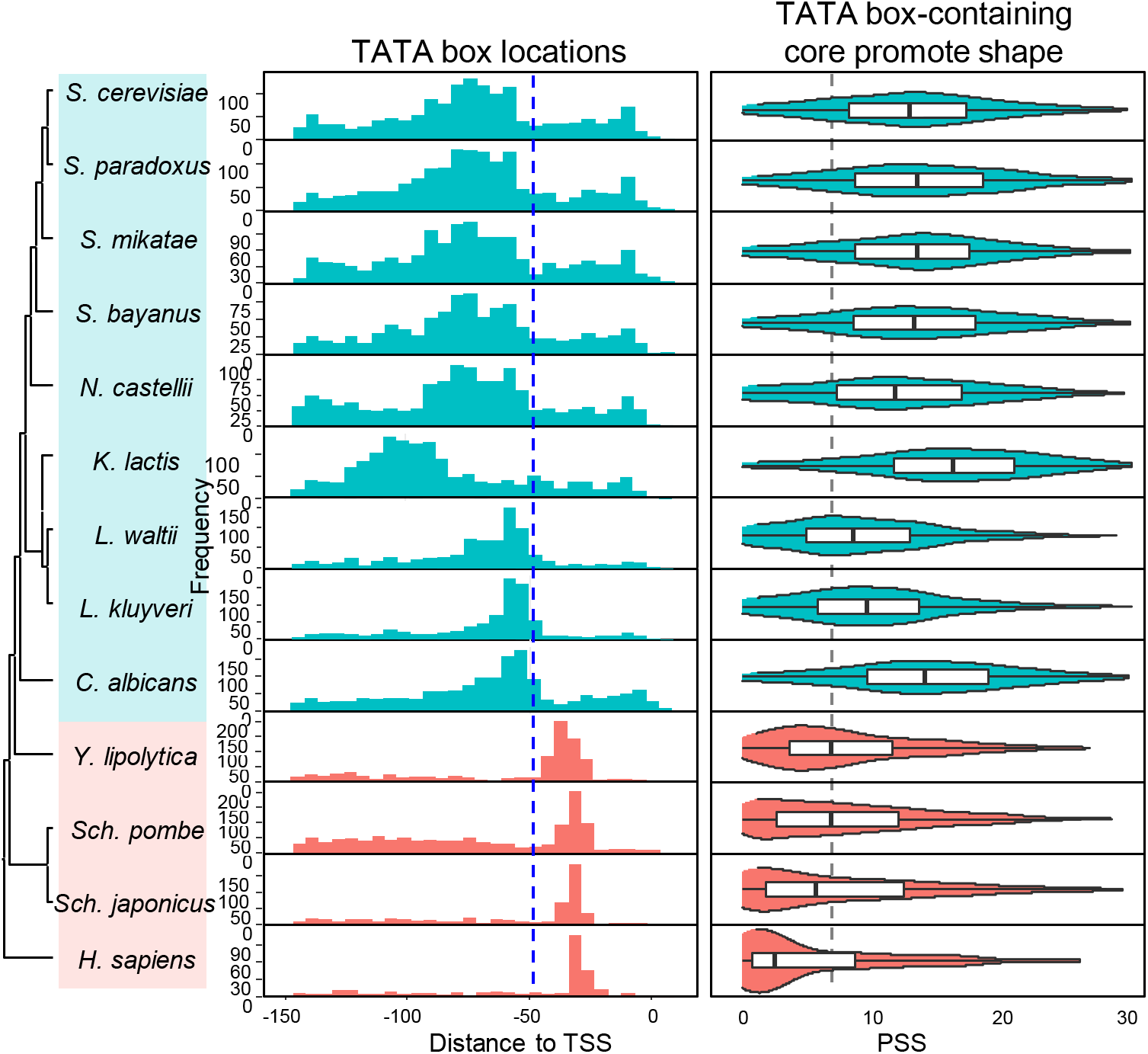
Evolution of transcription initiation mechanisms from TATA box-containing promoters. The left panel displays the phylogenetic relationships of the 12 yeast species using human as an outgroup. The middle panel shows the distributions of distances between TATA boxes and TSSs in each species. Blue dashed line refers to the position −50. The names of the “scanning model” species are shaded in cyan, and “classic model” species are shaded in orange. The right panel shows the promoter shape score (PSS) of TATA box-containing core promoters. The grey dashed line indicates the median PSS value in *Y. lipolytica*, *Sch. pombe*, and *Sch. japonicas*.

### Purine as the first recruited nucleotide is critical for accurate transcription initiation and efficient 5’capping

Our data show that a strong preference of PyPu dinucleotides at TSSs is present in all species examined (Fig. 3A; Supplemental Fig. S3). Unlike the pyrimidine at position −1, the functional role of purine at position +1 remains elusive. Unlike DNA replication, transcription initiation does not require an RNA primer, so the first NTP is added to the template strand by RNA pol II without forming a phosphodiester bond. Due to the difference between transcription initiation and extension, we speculated that the mismatch rate at the first position could be higher than its downstream sites. The strong preference of a single type of nucleotide for transcription initiation could theoretically reduce the probability of mismatch.

**Figure 3:**
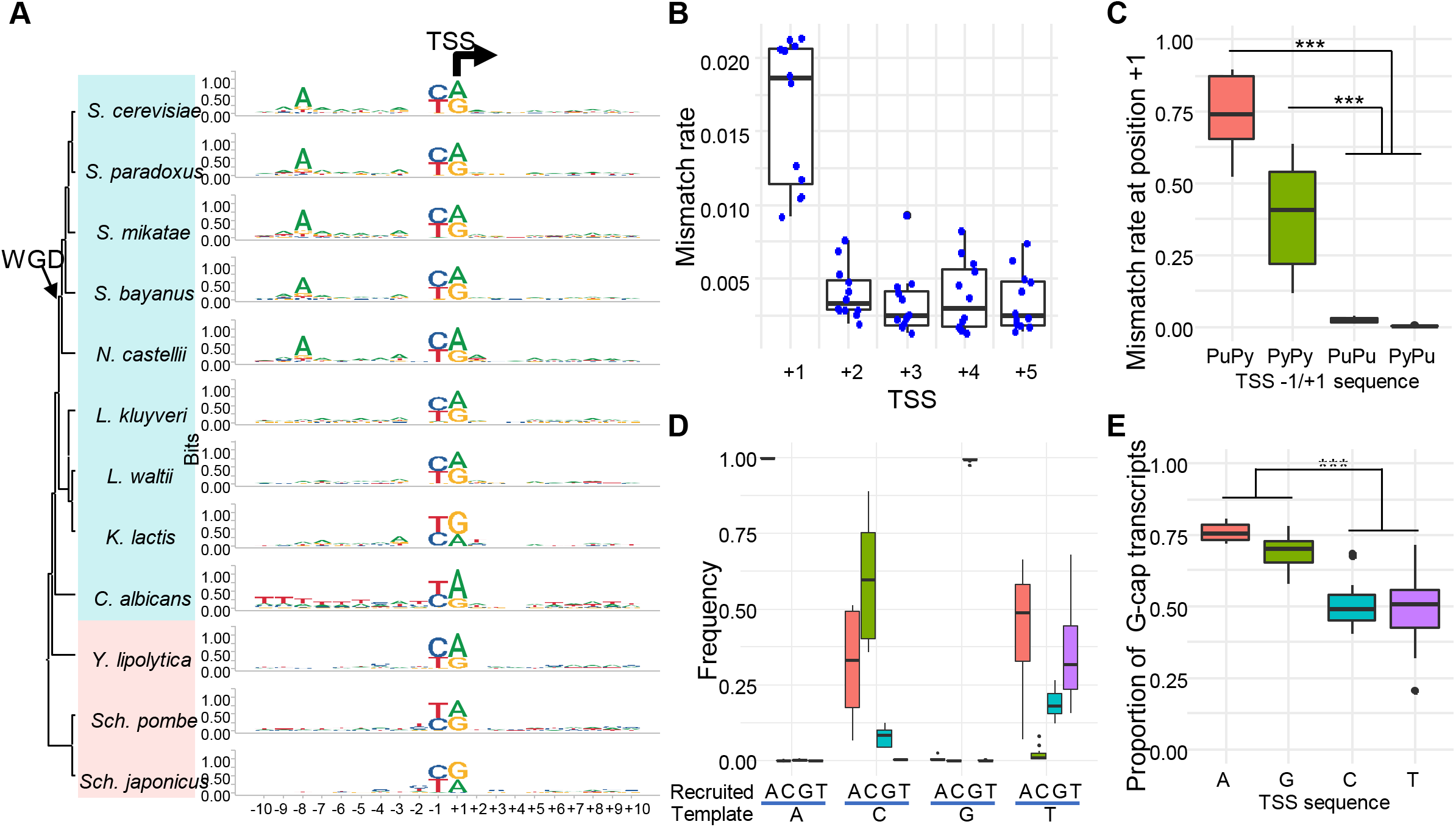
Functional roles of PyPu dinucleotide in transcription initiation. (A) The evolutionary changes of the consensus sequence of core promoters in the 12 yeast species. The black arrow indicates the TSS position and transcription direction. The arrow indicates the occurrence of WGD. (B) Boxplot of the distributions of mismatched rates at the first five sites of transcripts in the 12 species. Each blue dot represents the mismatch rate at each site in one species. (C) Mismatch rates in transcripts initiated from different TSS dinucleotides: PuPy, PyPy, PuPu, and PyPu. *: *p* < 0.01; **: *p* < 0.001; ***: *p* = 0. (D) Frequencies of each type of nucleotides at the first site of transcripts that are initiated from different nucleotides at the sense strand of genome. (E) Boxplot of the proportions of transcripts with a detected G-cap at the 5’end among transcripts with different types of nucleotides in the 12 yeast species.

The raw CAGE sequencing reads in the 12 yeast species allowed us to thoroughly test these hypotheses by comparing transcript sequences and their genomic templates (see Materials and Methods). As predicted, we found that the mismatch rates at the first position are ~ 6 times higher than that of the next four sites (Fig. 3B). We then examined the impacts of different dinucleotide combinations at +1/-1 position on initiation fidelity. The proportions of mismatched nucleotides in transcripts initiated from PyPu (0.003) and PuPu (0.026) are drastically lower than that from PyPy (0.405), and PuPy (0.740). This result demonstrates that purines at the +1 position of TSS are critical for recruiting the correct NTP to the first position, while the nucleotide at the −1 position has much less impact (Fig. 3C; Supplemental Fig. S4A). Interestingly, among those mismatches initiated from pyrimidine, we found that a purine, particularly adenine, is frequently incorporated by Pol II (Fig. 3D; Supplemental Fig. S4B). For instance, if the first nucleotide at the sense strand is thymine, an adenine is more likely to be recruited by Pol II than thymine, suggesting that purine is strongly preferred by Pol II at +1 position, regardless of template nucleotide.

The next question arises is that why Pol II prefers purine, especially adenine, as the initiation nucleotide. As a step of post-transcriptional modifications, a cap structure (e.g. N7-methylated guanosine, or m7G) is added to the first nucleotide of the primary mRNA, which is required for cap-dependent initiation of protein synthesis and prevention of exonuclease cleavage (Both et al. 1975; Muthukrishnan et al. 1975). To determine the impacts of different nucleotides at the 5’end of mRNA on 5’capping, we examined the raw CAGE sequencing reads to calculate the proportion of m7G caps for transcripts with each type of nucleotide at the 5’end. We found that transcripts with a purine at the 5’end have much higher rates of being capped by m7G (73.16%) than those with a pyrimidine (49.19%) (Fig. 3E; Supplemental Fig. S4C). Therefore, our results suggest that the strong preference of purine at the TSS provides the best chance for their transcripts being capped, increasing the probability of successful protein synthesis and reducing mRNA cleavage by exonuclease.

### The gain of an adenine-rich region immediately upstream of TSS during the evolution of “scanning model” yeasts might facilitate TSS selection

Adenine is present at eight bp upstream of most TSSs (−8A) in *S. cerevisiae* (Zhang and Dietrich 2005; Lu and Lin 2019). We found that the predominance of −8A also presents in other budding yeast species that have experienced an ancestral whole genome duplication (WGD) (Fig. 3A; Supplemental Fig. S5). By conducting a sliding window analysis of nucleotide frequency, we found an adenine-rich (A-rich) region that is present immediately upstream of TSS, with a peak at −8 position, in all “scanning model” yeasts but *Candida albicans* (Fig. 4A; Supplemental Fig. S4). As a human opportunistic pathogen, *C. albicans* is the earliest diverging lineage among the “scanning model” species. The TSS-proximity A-rich region is absent in all “classic model” species. Instead, they have an AT-rich region at ~30 bp upstream of TSS, corresponding to the location of the “TATA box” (Fig. 4A; Supplemental Fig. S5). Based on the phylogenetic distribution of the A-rich region, it is most parsimonious to infer that the enrichment of the TSS-proximity A-rich region had originated after the divergence of *C. albicans* during the evolution of budding yeasts. In addition, the common ancestor of WGD yeasts had gained a strong preference of adenine at a specific location (−8) within the A-rich region. The gain of −8A in WGD species suggests that there is a more stringent requirement of the distance between adenine and TSS for transcription initiation in the WGD species.

**Figure 4:**
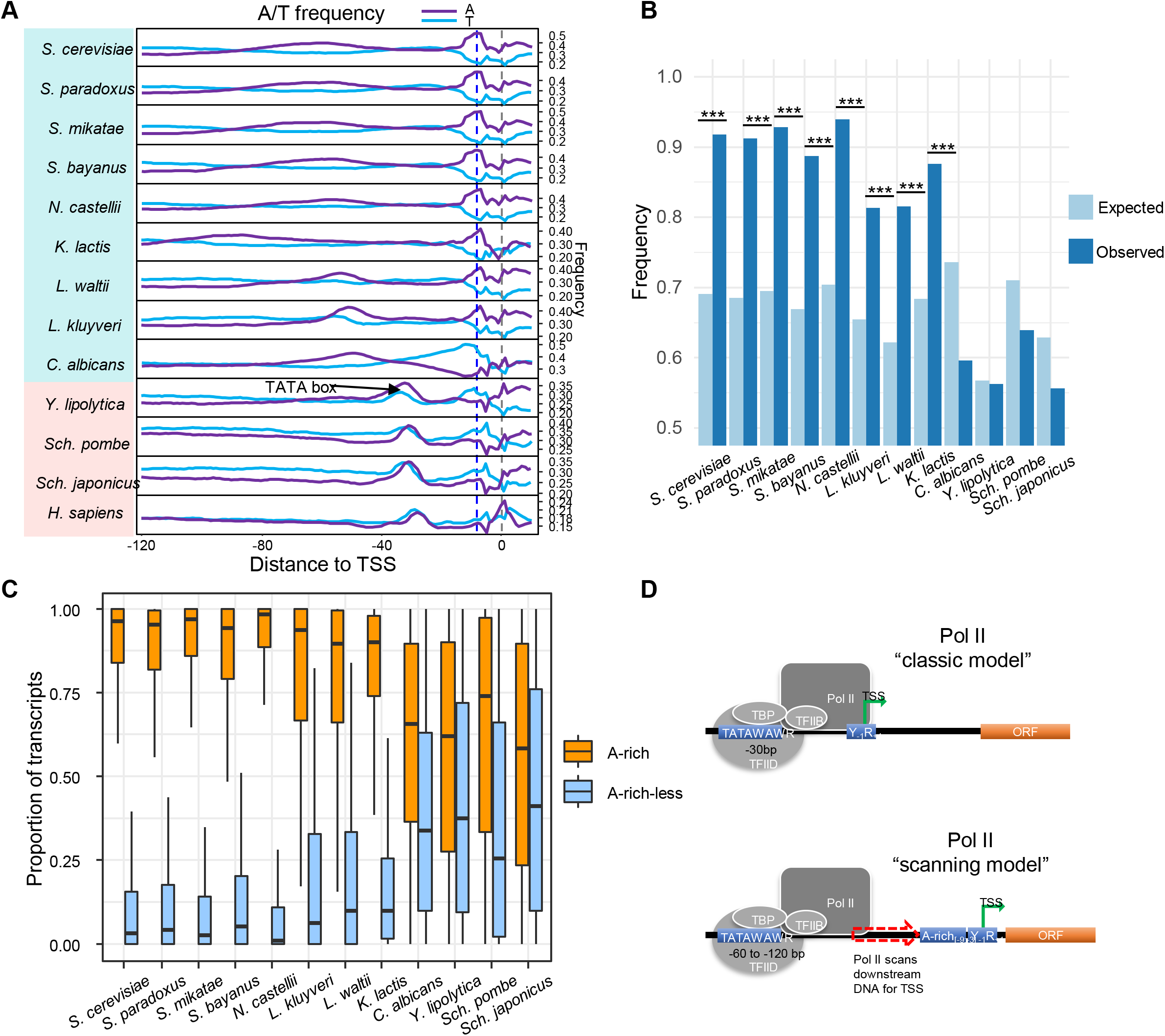
Presence of an adenine-rich region in the “scanning model” species and its functions. (A) A sliding window analysis of A/T frequencies. Window size is 6 bp with a step size of 1 bp. The black arrows indicate locations of the TATA box. Blue dashed line refers to the position 8 bp upstream of the TSS. The grey dashed line refers to the TSS position. (B) Frequency of expected and observed A-rich (at least 2 As) region in the window of 9 to 3 bp upstream of the TSS. ***: *p* = 0, Chi-square test. (C) Proportions of transcripts initiated from A-rich associated TSSs and TSSs without an A-rich region within core promoter regions in the 12 species. (D) A comparison of the “classic model” and “scanning model” of transcription initiation. We propose that in the “scanning model”, the A-rich region functions as a binding motif to interact with a component of PIC, which stabilizes the PIC and Pol II and facilitates efficient transcription initiation from its downstream PyPu sites.

Structural studies showed that the −8A in *S. cerevisiae* is recognized by the B-reader helix of TFIIB, which is required for TSS selection (Kostrewa et al. 2009; Sainsbury et al. 2013). It is reasonable to postulate that adenines in the A-rich region serve as a binding motif for a PIC component, probably TFIIB, in the “scanning model” species. The interaction between TFIIB and the A-rich region might temporarily pause the scanning process, and direct Pol II to initiate transcription from its downstream PyPu. Therefore, the A-rich region in “scanning model” species might have a similar role as the TATA box in the “classic model”, serving as an anchor point for PIC to initiate transcription.

If the A-rich region has its functional importance, it should be overrepresented immediately upstream of TSS in “scanning model” species, and such enrichment should be absent in the “classic model” species. Here, we defined an A-rich region as a minimal number of 2 As between −9 to −3 nt upstream of TSS (Supplemental Fig. S6). Based on this definition, we observed that the A-rich region is present in 91.80% of TSSs in *S. cerevisiae*, comparing to the expected value of 69.1% based on nucleotide frequencies (*p* = 1.8×10^-227^, Chi-squared test, Fig. 4B). Such enrichment is found in all “scanning model” species, excluding *C. albicans*. In contrast, the A-rich region is not enriched or underrepresented in all “classic model” species (Fig. 4B). Considering that one PyPu dinucleotide is expected to be found in every window of 5 bp by chance, it also explains why TSSs are located a few bp downstream of the A-rich regions in “scanning model” species.

Transcription initiation from a core promoter may occur from an array of nearby TSSs, and some TSSs may lack an upstream A-rich region. If the A-rich region is required for efficient transcription initiation in the “scanning model” species, we expected that most transcript s within a core promoter should be initiated from the TSSs associated with an A-rich region. As shown in Fig. 4C, we found that the proportion of transcripts initiated from A-rich associated TSSs are much higher than A-rich-less TSSs in all “scanning model” species but *C. albiacans*, supporting our hypothesis. In *C. albicans,* we observed a “thymine-rich” region upstream of its TSSs (Supplemental Fig. S5), suggesting that the molecular mechanisms of transcription initiation in *C. albicans* might be different from other “scanning model” species. Based on these findings, we proposed a model of transcription initiation in the “scanning model” species, in which the A-rich region serves as an anchor point for PIC. If a PyPu is available within 3-9 bp downstream of the A-rich region, the PIC recruits Pol II to starts transcription initiation. Otherwise, the PIC continues to scan the promoter sequence until it reaches to the favorable sequence combination (Fig. 4D).

### Genetic basis underlying evolutionary conservation and divergence of core promoters

Our results suggest important roles of PyPu and the A-rich region (or −8A in WGD species) in transcription initiation. We then aimed to examine the mutation patterns of these sequence elements to better understand their impacts on the evolutionary divergence of TSSs and core promoters. We focused on three closely related species, *S. cerevisiae, S. paradoxus* and *S. mikatae,* which allowed us to align the entire genomes for accurate identification of orthologous core promoters. Using *S. mikatae* as an outgroup, we divided orthologous core promoters into three groups: “Conserved”, “Shifted” and “Turnover (gain/loss)” (see Materials and Methods, Fig. 5A). The “Conserved” group includes the orthologous core promoters that are present in both species, and the positions of dominant TSSs remain the same. If the dominant TSS has changed in one or both species, it was defined as the “Shifted” group. If a core promoter is lost or newly gain in *S. cerevisiae* or *S. paradoxus,* it was classified as the “Turnover” group.

**Figure 5:**
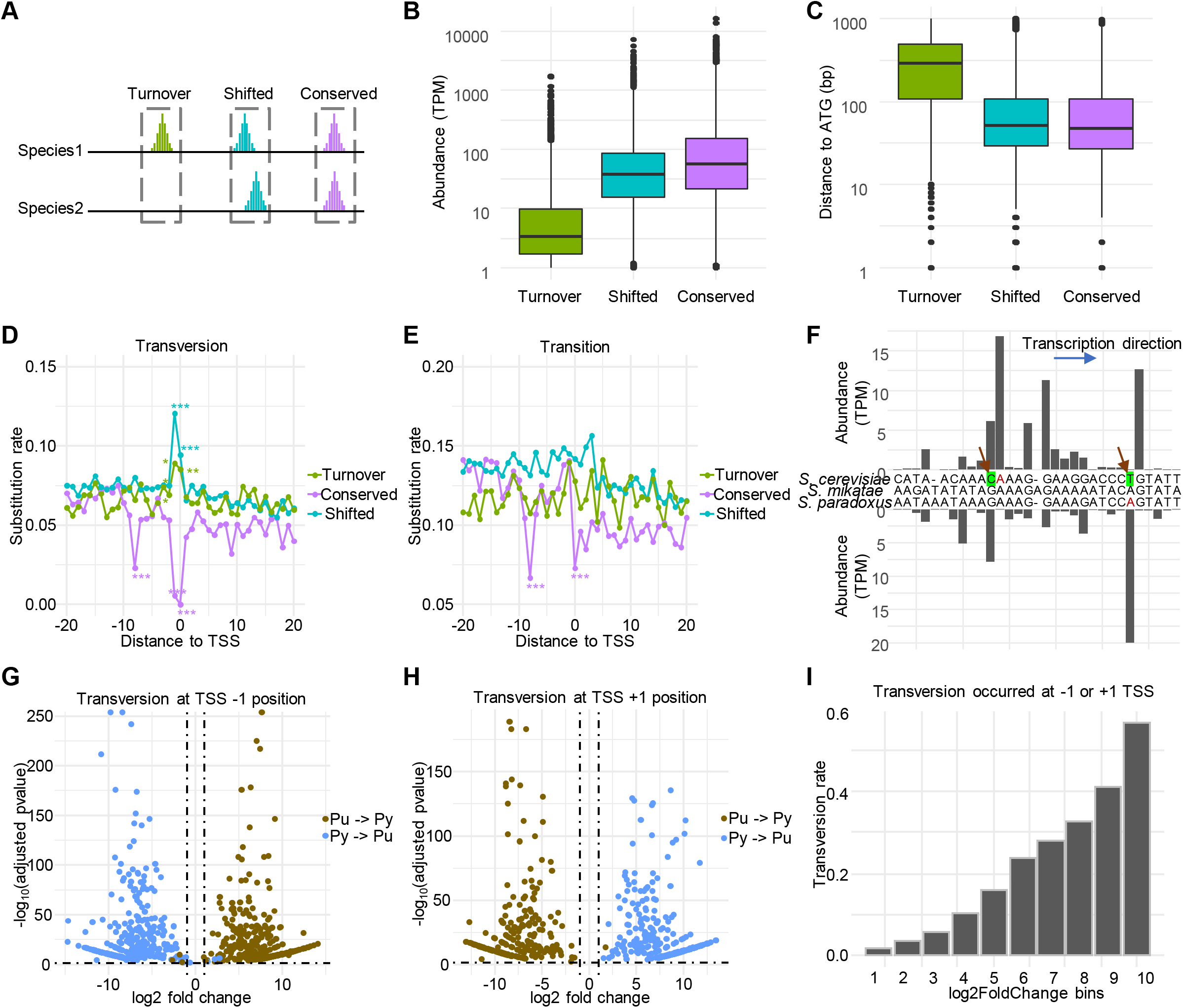
Genetic basis underlying the evolutionary divergence of core promoters and TSSs in budding yeasts. (A) A schematic diagram of three types of core promoters identified based on their evolutionary patterns. (B) Different transcriptional activities among the three types of core promoters. (C) The “Turnover” core promoters tend to locate more upstream from the ATG codon than the other groups. (D) The rate of transversional substitution at each site in a 41-bp region surrounding the TSS (−20 to +20), which were calculated between orthologous core promoters in *S. cerevisiae* and *S. paradoxus.* The sites with significantly higher or lower substitution rates are indicated by * (Chi-squared test). *: *p* < 0.01; **: *p* < 0.001; ***: *p* = 0. (E) The rate of transitional substitution at each site in a 41-bp region surrounding the TSS ( −20 to +20). (F) Example of a “Shift” type core promoter and its associated genomic sequences. This core promoter was assigned to the genes in the orthologous group of *YDL182W.* New mutations in *S. cerevisiae* and *S. paradoxus* are indicated by arrows. (G) Volcano plot illustrates that mutations from pyrimidine to purine (Py→Pu) and from purine to pyrimidine (Pu→Py) at −1 TSS position have the opposite impacts on transcriptional activities of TSSs. The horizontal dashed line refers to the adjusted p-value of 0.05, and the vertical dashed line refers ≥ 1 log_2_ fold change. (H) Volcano plot illustrates that mutations from pyrimidine to purine (Py→Pu) and from purine to pyrimidine (Pu→Py) at +1 TSS position have the opposite impacts on transcriptional activities of TSSs. (I) The TSSs with larger fold changes in transcriptional activities are more likely to be associated with transversional mutations at position −1/+1.

Overall, we found that core promoter turnovers are prevalent. *S. cerevisiae* had gained 670 and lost 229 core promoters since its divergence from *S. paradoxus,* accounting for 10.3% and 3.5% of all its core promoters (Supplemental Fig. S7A). Similar patterns are present in *S. paradoxus.* Compared with conserved and shifted core promoters, the newly gained or lost core promoters tend to have lower transcriptional activities (Fig. 5B) and are usually located at further upstream of the translation start codons (Fig. 5C).

We examined the genomic sequences from −20 to +20 nt surrounding the dominant TSSs for each group of core promoters to infer their associated genetic changes. We observed distinct patterns of genetic divergence at −1, +1, and −8 positions among the three types of core promoters. In the “Conserved” type, the rates of nucleotide substitutions, particularly transversions, are nearly depleted at −1, +1, and −8 positions (Fig. 5D-E), suggesting that the nucleotide type at these positions is critical for maintaining core promoter activities. In contrast, elevated rates of transversional substitutions are observed at −1, +1 positions in the “Shift” and “Turnover” groups (*p* < 0.001, Chi-square test) (Fig. 5D). For example, the core promoter of *YDL182W* in *S. cerevisiae* has experienced a change of its dominant TSS since its divergence from *S. paradoxus* (Fig. 5F). The ancestral dominant TSS was lost in *S. cerevisiae* due to a transversion mutation that replaced adenine with thymine at −1 position, converting PyPu to PyPy. Meanwhile, *S. cerevisiae* has gained a new dominant TSS that is 13 nt upstream of the ancestral one by replacing guanine to cytosine, generating a new PyPu dinucleotide (Fig. 5F). Besides, in the “Turnover” group, the active core promoters have a significantly higher frequency of PyPu and −8A than their counterparts (silent core promoters) (Supplemental Fig. S7B). However, the frequency of the TATA box is similar between the two groups, supporting that nucleotide turnovers at −1, +1 and −8 positions play an important role in the evolutionary divergence of core promoter activities.

We then evaluated the impacts of transversions at −1 and +1 positions on transcription initiation activities. If the nucleotide at position −1 changes from the preferred pyrimidine to purine, most of them are associated with significantly reduced transcriptional activities, and an opposite pattern was observed if it is changed from purine to pyrimidine (Fig. 5G). At position +1, because purine is the preferred nucleotide, a change from a purine to pyrimidine significantly reduced its transcriptional activities, or vice versa (Fig. 5G). These results further support the importance of PyPu dinucleotide in transcription initiation. By dividing TSSs based on the number of fold changes between the two species, we found that the proportion of TSSs with transversion mutations at +1/-1 positions increases as the fold changes in transcriptional activities increase. In the group of TSSs with the largest fold changes, 56.7% of them are associated with transversion mutations at +1/-1 positions (Fig. 5I), suggesting that the TSSs with the most significant evolutionary divergence in transcriptional activities are more likely due to transversional mutations at +1/-1 positions.

### Genetic determinants of core promoter shape

We observed that in the “classic model” species, TATA box-containing core promoters tend to have a sharper core promoter than TATA-less ones (Supplemental Fig. S2), suggesting an impact of TATA box on core promoter shape. However, within each type of core promoter, there are large variations in core promoter shape. In addition, in the “scanning model” species, we did not observe a significant difference in core promoter shape between them. Therefore, the major determinants of core promoter shape require further investigations. Our previous study showed that the number of TSSs, the spacing of TSS, and distribution transcription activities among TSS determines the shape of a core promoter (Lu and Lin 2019). Here, we found that the number of TSSs within a core promoter could explain 85% of the variance of core promoter shape (R^2^ = 0.85, Fig. 6A).

**Figure 6:**
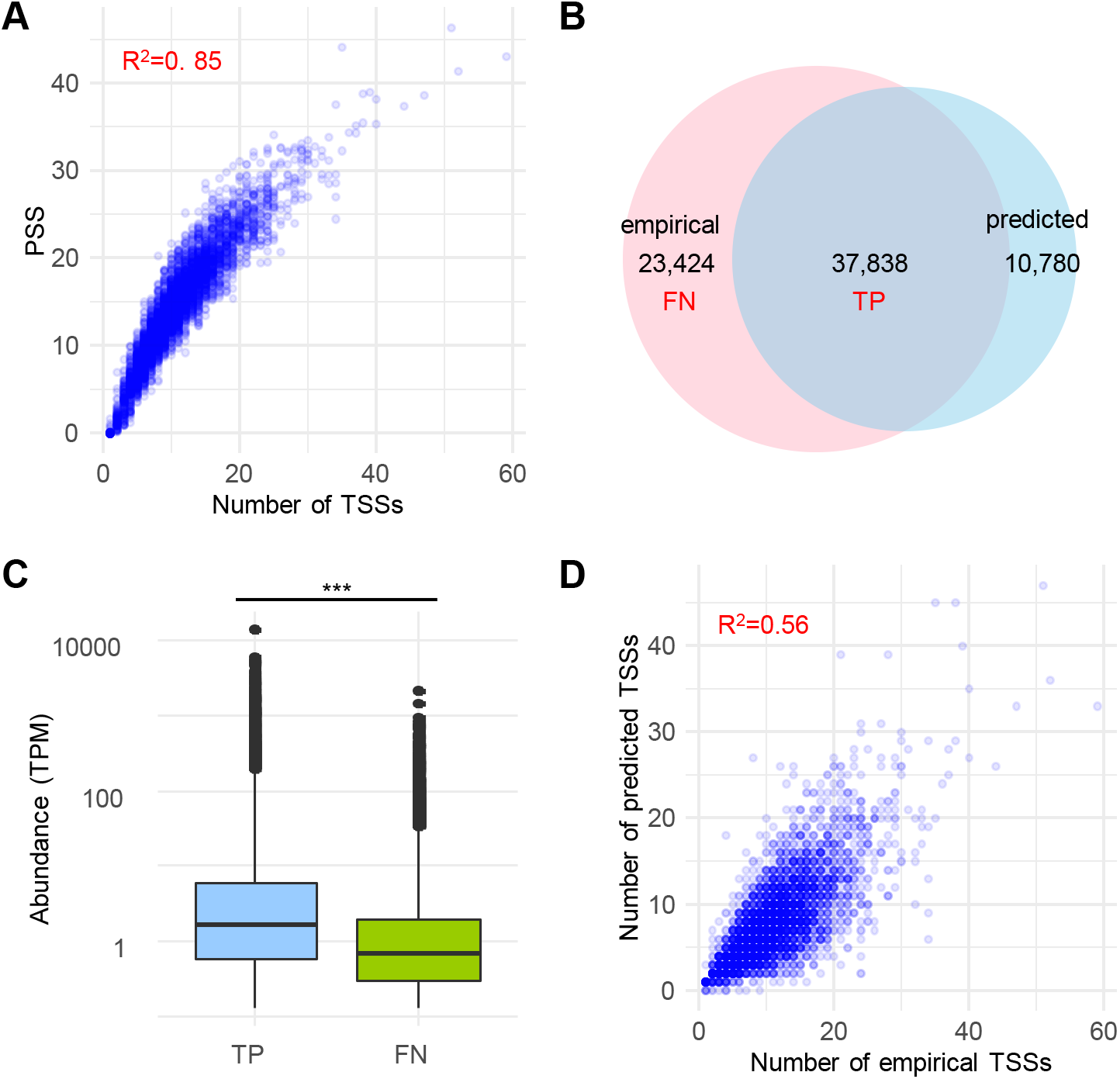
The genetic determinants of core promoter shape. (A) Scatter plot showing a positive correlation between the number of observed TSSs and core promoter shape score (PSS). (B) Venn diagram showing the number of TSSs identified by CAGE and TSSs predicted from the core promoter sequences. (C) Boxplot showing a higher transcription activity in truepositive (TP) TSSs than in false-negative (FN) TSSs. (D) Scatter plot showing a positive correlation between the number of observed TSSs and predicted TSSs in each core promoter.

Our sequence analysis suggested that the number of TSSs is likely determined by the sequence feature in a core promoter, because the PyPu and A-rich region are present in most of TSSs. To further test this hypothesis, we used the presence of A-rich (at least 2 As) at positions −9~-3 and PyPu dinucleotides at positions −1,+1 to predict TSSs within a region of − 30,+30 around the dominant TSSs of core promoters in the “scanning model” species *S. cerevisiae*. Our prediction method successfully predicted most TSSs observed by CAGE, with a true positive rate (TPR) of 61.8% (Fig. 6B). Our comparison of transcription activities of TSSs between true positives (TP) and false negatives (FN) showed that FN TSSs have significantly lower transcriptional activities (Fig. 6C). TSSs observed by CAGE could be due to stochastic transcription near a strong TSS or technical artifacts, which tend to have lower transcriptional activities than canonical TSSs. We speculated that many of these FN TSSs could belong to those non-canonical TSSs. Moreover, we observed a strong positive correlation between the number of TSSs predicted by sequence features and the number of TSSs observed by CAGE (R^2^ = 0.56, Fig. 6D). These observations indicate that the intrinsic sequence features of a core promoter largely determine its core promoter shape.

### Other common motifs in yeast core promoters

A GA element (GAAAAA) has been identified as a conserved promoter element in TATA-less promoters in *S. cerevisiae* (Seizl et al. 2011). Interestingly, we found that the GA element is enriched in core promoter regions in all “scanning model” yeast species, and they are located at similar positions with the TATA box (Supplemental Fig. S8). These findings suggest that the GA element might function as binding sites for general transcription factors in the “scanning model” species, supporting the presence of two distinct Pol II transcription initiation machinery in yeasts.

Our TSS maps allowed us to predict putative core promoter motifs in yeasts by using *de novo* motif discovery methods. Here, we showed that, besides the TATA box, eight motifs are significantly enriched in the core promoter regions in at least three yeast species (Supplemental Fig. S9A). These shared motifs are generally located at similar locations relative to the TSS within each type of transcription initiation mechanism (Supplemental Fig. S9B), further supporting the presence of two models of Pol II initiation. For example, the motifs REB1, ABF1, and TOD6, which were found in most “scanning model” species, are located at similar positions of TATA box. These motifs, as well as the GA element, have a small portion of co-occurrence with the TATA box (Supplemental Fig. S10), indicating that these motifs might serve as alternative binding sites of GTFs.

## DISCUSSIONS

In this study, we generated quantitative TSS maps for ten budding yeasts and two fission yeasts, representing the most comprehensive TSS atlas in yeasts so far. These TSS maps improve genome annotations for these species by providing 5’ boundaries for most protein-coding genes at a single-nucleotide resolution. Most importantly, our comparative studies of these TSS maps revealed the origin of the “scanning model” of transcription initiation, and sheds light on the molecular mechanisms of transcription initiation by inferring the functional roles of PyPu and A-rich region in core promoters.

### The origin and stepwise evolution of the “scanning model” in budding yeasts

One of the most significant findings of this study is that the shift of transcription initiation from the “classic model” to the “scanning model” occurred after the split of *Y. lipolytica* during the evolution of budding yeasts. Our study indicates that the transition from the “classic model” to the “scanning model” is likely a stepwise process that involved multiple genetic innovations in both PIC genes and promoter sequences that occurred at different evolutionary stages.

The first major evolutionary event is the switch of the TSS selection process in the ancestral budding yeast after its splitting from the *Y. lipolytica* lineage. It could be due to genetic innovations in GTFs, such as TFIIB, and PoI II, resulting in a distinct transcription initiation machinery. By swapping TFIIB and Pol II from *S. cerevisiae* to *Sch. pombe*, it was found that *Sch. pombe* transcription initiated 40-120bp downstream of the TATA box, suggesting that TFIIB and Pol II play a key role in determining the TSS (Li et al. 1994). Another factor could be the divergence of nucleosome occupancy patterns in promoter regions. It was found that *S. cerevisiae* has a wider nucleosome depletion region (NDR) immediately upstream of TSS than *Sch. pombe* (Moyle-Heyrman et al. 2013). We observed a similar pattern for the group of TATA box-containing core promoters between the two species based on published nucleosome occupancy data (Brogaard et al. 2012; Moyle-Heyrman et al. 2013) (Supplemental Fig. S11). Therefore, the wider NDR in *S. cerevisiae* provides a longer naked DNA that could facilitate the scanning process. Comparative studies of sequences of each PIC component would be needed to infer other critical genetic changes associated with the origin of the “scanning model”.

The second evolutionary event is the gaining of the A-rich region in ancestral budding yeasts after their divergence from *C. albicans*. In the “scanning model”, a different mechanism of TSS selection should be involved for the PIC to pause the scanning process and initiate transcription. Here, we showed that an A-rich region exists immediately upstream of TSSs in all “scanning model” species after their divergence from *C. albicans*. In *S. cerevisiae*, TFIIB interacts with adenine at the −8 position (thymine at the template strand) in the A-rich region to facilitate TSS selection and transcription initiation (Kostrewa et al. 2009; Sainsbury et al. 2013). We propose two possible functional roles of the A-rich region in the “scanning model” species. First, the A-rich region might stabilize the PIC through its interaction of one of its components, such as TFIIB, increasing transcription efficiency. Second, it could reduce the number of usable TSSs to decrease the production of undesired transcript isoforms. Because one PyPu is expected to be found in each window of 5 nt, a requirement of presence of A-rich region immediately upstream of PyPu largely eliminates initiation from other PyPu sites. However, how PIC components interact with the A-rich region interact requires further studies.

The most recent evolutionary event is the origin of the preference of −8A in the A-rich region in the WGD species. Our results show that the −8 position is nearly depleted of any types of substitutions in the group of conserved core promoters (Fig. 5D-E). This is consistent with a previous study that mutation of −8A in *S. cerevisiae* led to almost complete loss of its corresponding transcription activity (Kostrewa et al. 2009). These observations support an important role of −8A in the “scanning model” of transcription initiation. The positional preference of adenine in WGD species might be due to the divergence in a PIC component that directly interacts with adenines. Comparative analyses of sequence and structural features for each PIC component between the WGD and non-WGD species will be necessary to better understand the function of the A-rich region and genetic mechanism underlying the changes of positional preference of adenine in the A-rich region.

### The important role of PyPu in transcription initiation

Another major finding of our study is that it improves the understanding of the role of PyPu during transcription initiation. The strong preference of purine as the first base of transcripts is likely due to the intrinsic preference of Pol II, which benefits subsequent post-transcriptional modification and protein synthesis. We found that transversions at position −1, and/or +1, which disrupt PyPu dinucleotide, results in remarkable changes of TSS activities and TSS shift, supporting their importance in transcription initiation. Furthermore, these results uncover a key genetic mechanism underlying the evolutionary divergence of TSSs and core promoters. We found that disruption of the PyPu sites is sufficient to eliminate its transcriptional initiation activities (Fig. 5D). However, other factors should play a more important role in gaining a new TSS. In most cases, the birth of a new TSS in a promoter region does not require mutations to obtain a PyPu dinucleotide due to its prevalence (one PyPu in every five bp). A study in the human genome found that most new TSSs emerged from transposable elements due to retrotransposon activities (Li et al. 2018). However, yeasts were known for scarce of active transposable elements (Bleykasten-Grosshans and Neuveglise 2011). We speculate that evolutionary innovations in *trans*-or *cis*-regulatory factors probably play a more important role in the birth of new TSS in yeasts.

### Core promoter shape is intrinsically determined by its sequence features

Core promoter shape has been considered as a genetically controlled molecular trait that is conserved in mammalian cell lines (Carninci et al. 2006), between development stages in *Drosophila* (Hoskins et al. 2011), and between mouse and human (Carninci et al. 2006). And yet, how genetic basis determines core promoter shape is not fully understood. Here we showed that core promoter shape is significantly correlated with its number of TSSs, which is intrinsically determined by sequence features, including A-rich at −9 to −3 positions and PyPu at − 1, +1 positions in *S. cerevisiae.* In the TATA box-containing genes, the TATA box serves as a significant landmark for the locations of core promoters. Such structure is valuable for the prediction of TSS and core promoters in the TATA box-containing genes, even though the distances between TATA box and core promoters are different between the “classic model” and “scanning model’ species. However, TATA box only accounts for a small portion of protein-coding genes (Basehoar et al. 2004; Yang et al. 2007). A better understanding of other binding sites of general transcription factors would be necessary for accurate prediction of core promoters and TSSs at a genome scale.

## MATERIALS AND METHODS

### Yeast strains and CAGE sequencing

Twelve yeast species, including ten budding yeast species and two fission yeast species, were used in this study (Supplemental Table S1). Strains were grown to log-phase in rich media (YPD liquid culture) at 30 °C. 5ug of total RNA was extracted with TRIzol (Invitrogen) from each sample. Two biological replicates of sequencing libraries were constructed for each yeast species following the nAnT-iCAGE protocol (Murata et al. 2014), and each nAnT-iCAGE library was sequenced using Illumina NextSeq (single-end, 75-bp reads) by the DNAFORM, Yokohama, Japan. Human CAGE sequencing data was downloaded from Adiconis *et al.* (Adiconis et al. 2018) and reanalyzed in this study following the same criteria.

### Inference of phylogenetic relationships and divergence times for the 12 yeast species

The phylogenetic relationships of the 12 species were inferred by the Maximum Likelihood method based on the largest subunit of Pol II RPB2 protein sequences. The divergence times between every two species were estimated by the RelTime method (Tamura et al. 2012), which were calibrated by the estimated divergence times obtained from TimeTree (Kumar et al. 2017).

### TSS calling and identification of core promoters based on CAGE data

The sequenced tags were mapped to each respective reference genome (Supplemental Table S1) using HISAT2 (Kim et al. 2015) with ‘--no-softclip’ option to avoid false TSSs. The reads mapped to rRNA sequences (28S, 18S, 5.8S, and 5S) were identified by rRNAdust (http://fantom.gsc.riken.jp/5/sstar/Protocols:rRNAdust) which changes the FLAG column in SAM files as ‘not passing filters’. The modified SAM files were then converted into BAM format and sorted by SAMtools (Li et al. 2009) for subsequent TSSs calling. Replicates of CAGE tags supporting each TSS from each yeast species were counted and merged. TSSs were omitted when their number of supporting CAGE tags are significantly smaller than the expected numbers according to Poisson Distribution. The transcriptional activity of each TSS was calculated as tags per million of uniquely mapped tags (TPM). TSSs in an approximate region are likely regulated by the same set of promoter elements and give rise to a functionally equivalent set of transcripts, which can be grouped into a single tag cluster (TC), representing a putative core promoter. We developed a “Peakclu algorithm” to identify TCs in each species. Briefly, we first identified a CAGE peak within a window of a minimum of 100bp. The surrounding TSSs were grouped with the CAGE peak into a TC. For each TC, we calculated the positions of 10^th^ and 90^th^ percentile based on a cumulative distribution of CAGE tags. TCs were assigned to Pol II transcribed genes using the same criteria as previously described (Lu and Lin 2019).

### Calculation of 5’UTR length and core promoter shape

Transcription is usually initiated from an array of TSSs, instead of a single TSS, and gives rise to a set of functionally equivalent transcripts with slightly different lengths of 5’UTRs (Kodzius et al. 2006). In many cases, multiple core promoters are concurrently used that generate significantly different lengths of 5’UTRs (Lu and Lin 2019). Using one TSS to calculate 5’UTR length cannot accurately represent the uncertainty and complexity of transcription initiation. We calculated the weighted 5’UTR length for each gene, which is the average 5’UTR length in all transcripts generated by a gene:

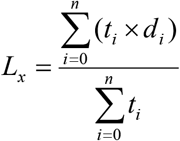

where *n* is the total number of TSSs identified for gene *X*, *t_i_* is the number of CAGE tags mapped to the *i*^th^ TSS, and *d_i_* is the length of 5’UTR in transcripts generated from the *i*^th^ TSS.

Core promoter shape score is defined based on the PSS formula (Lu and Lin 2019) and modified as:

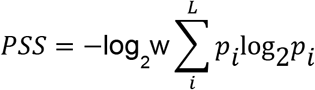

where *p* is the probability of observing a TSS at *i*^th^ TSS within a core promoter, *L* is the total number of TSSs that pass filtrations by Poisson Distribution, and *w* is the core promoter width which is defined as the distance between 10^th^ and 90^th^ quantiles.

### Analysis of consensus transcription initiation sequence

Sequences (±10 nt) surrounding the dominant TSS of all core promoters from each yeast species were extracted. Sequence motifs were plotted with seqLogo package (Bembom 2019) in R. The frequency of mismatched nucleotide at the +1 TSS position was calculated with G-capped reads only. To mitigate transcriptional or technical noise in coding regions, we only considered tags whose first nucleotide mapped in the core promoter region from quantile 0.1 to 0.9 to calculate G cap rates.

### Analysis of orthologous core promoters

Orthologous core promoter analyses were conducted among *S. cerevisiae, S. paradoxus*, and *S. mikatae*. Pairwise genome alignments were carried out with wgVISTA (Frazer et al. 2004). To minimize background noise, only the sequences of core promoters with TPM >= 1 were used as queries to search for their orthologous core promoters. Orthologous core promoter groups were later discarded if they are not associated with protein-coding genes (Supplemental Dataset S15). If a “Turnover” core promoter was generated by insertion and deletions, it was excluded in subsequent analysis analyses.

### Analysis of TATA box and *de novo* motif discovery

To identify TATA box, we first generated a TATA box matrix based on a consensus sequence of TATAWAWR by seq2profile.pl in the HOMER package (Heinz et al. 2010) with zero mismatches allowed. We then used the generated TATA box matrix to search against genomic sequences for occurrence and locations of TATA box in each yeast species using findMotifs.pl in HOMER. To identify novel sequence motifs enriched in the promoter regions, we performed *de novo* motif discovery for sequences that are from 100 bp upstream to 50 bp downstream of the dominant TSS in each core promoter by HOMER. The occurrence and locations of the predicted motifs from each species were identified using findMotifs.pl.

### Data access

The raw sequencing data generated in this study have been submitted to the NCBI BioProject database under accession number PRJNA510689. The quantitative maps of TSSs and core promoters generated in this study can be visualized and downloaded from the YeasTSS database (http://www.yeastss.org) (McMillan et al. 2019).

## ACKNOWLEDGEMENTS

This study was supported by the President’s Research Fund from Saint Louis University and U.S. National Science Foundation (NSF 1951332) to ZL. We would like to thank Dr. Genevieve Fourel for constructive comments.

